# Assessing behavioral sex differences to chemical cues of predation risk while provisioning nestlings in a hole-nesting bird

**DOI:** 10.1101/2022.03.14.482199

**Authors:** Irene Saavedra, Gustavo Tomás, Luisa Amo

**Affiliations:** Departamento de Ecología Evolutiva, Museo Nacional de Ciencias Naturales (CSIC), C/ José Gutiérrez Abascal, 2. E-28006, Madrid, Spain; Departamento de Ecología Funcional y Evolutiva, Estación Experimental de Zonas Áridas (EEZA-CSIC), Carretera de Sacramento, s/n. La Cañada de San Urbano, E-04120, Almería, Spain; Departamento de Ecología Evolutiva, Universidad Rey Juan Carlos. C/ Tulipán, s/n. 28933, Madrid, Spain

**Keywords:** anti-predator behavior, chemical cues, olfaction, perceived predation risk, risk taking, sexual differences.

## Abstract

Birds can assess nest predation risk and adjust their parental activity accordingly. Risk taking behavior should be related to investment in reproduction as well as to confidence in parenthood that often differ between sexes. In those cases, sexual differences in risk taking behavior may be expected. For example, in blue tits, females invest more time and energy than males in nest-building, egg laying and incubation. Furthermore, confidence in parenthood is supposed to be higher for females, as extrapair paternity is common in this species. Therefore, the reproductive value of nestlings may be higher for females than for males and the former may assume greater risks to ensure nestling growth and maximize their reproductive success. We examined potential sexual differences in the risk assumed by parents in relation to perceived risk of predation inside the nest cavity, where predation risk perception may be higher. We increased perceived predation risk by adding predator chemical cues inside blue tit (*Cyanistes caeruleus*) nest-boxes, and we tested whether female and male parents differed in the risk assumed when taking care of nestlings. Females and males did not differ in the risk assumed in response to perceived predation risk. However, females reduced time devoted to nest sanitation activities when predator chemical cues were detected inside the nest-box, likely as an anti-predatory strategy to minimize their own risk of predation. Therefore, these results add to the evidence that birds can detect chemical cues of predators inside the nest cavity and suggest that the behavioral response to an increase in risk of predation perceived through olfactory cues is not sex-dependent in blue tit.

## Introduction

Parental care enhances the fitness of offspring [1] but such parental activity during reproduction can reveal the location of the nest to predators. Predators typically locate nests during nest building, incubation recesses and provisioning visits to nestlings [2]. Thus, an important cost of parental care comes from an increase in the risk of nest predation [3,4] and in the probability of mortality [1]. Parents face a trade-off between maintaining an appropriate level of parental care and reducing predation risk, both for themselves and their offspring [3,4]. Therefore, evolution has favored different parental behavioral strategies to reduce the risk of predation while maximizing reproductive success [5–7].

Several studies have shown that birds can assess nest predation risk and adjust their parental activity accordingly [8–14]. Many bird species reduce their parental activity under a perceived increase in predation risk [4,10,15–18]. For example, blue tits (*Cyanistes caeruleus*) hesitate to enter the nest-box, delaying their entry, and reduce the time spent inside the nest-box, when nests contain predator scent [11]. Blue tits also reduce the time spent inside the nest-box in non-essential activities for nestling survival, such as nest sanitation, when exposed to either visual or chemical cues of a predator in the vicinity of their nest [14]. During nest sanitation activities, blue tits remove ectoparasitic arthropods (e.g. mites, blowfly larvae, fleas) from nestlings or nest material. Therefore, this antipredatory behavior may entail indirect costs in terms of parental and nestling fitness [19]. However, they did not modify provisioning rates to ensure nestling growth [11,14]. In contrast, other bird species reduce provisioning rates to minimize detectability by predators when they perceive a high risk of predation [4, 13, 20–23]. For example, Siberian jays (*Perisoreus infaustus*) modify their daily nest visitation patterns in territories with a higher level of predation risk [8]. Orange-crowned warblers (*Vermivora celata*) also reduce provisioning rates in nests poorly concealed to predators [12]. In contrast, pied flycatcher (*Ficedula hypoleuca*) parents [24] and European roller (*Coracias garrulus*) males [25] increase their nestling provisioning rates when exposed to an increase in the perceived risk of predation, probably to silence the begging calls of nestlings to avoid drawing attention to the nest location, or to boost nestling growth to advance fledging time [24,26]. The variability in strategies exhibited by birds for minimizing predation during reproduction may greatly depend on the type of predator. Parents usually increase nest visitation in response to a nest predator such as a snake to silence offspring (e.g., [25]) while they decrease nest visitation after exposure to an aerial predator because of the danger to themselves (e.g., [27]; but see [28]).

Parental care and parental investment differ between females and males in numerous taxa, from invertebrates to vertebrates, including fishes, birds and mammals [1,29,30]. In socially monogamous birds, although both parents usually take care of their offspring, investment in parental care often differs between females and males [31]. For example, blue tit females alone build the nest, lay the eggs, and incubate them [32]. Female blue tits also spend more time at the nest and visit the nest more times to feed the nestlings than males [33]. Also, tree swallow (*Tachycineta bicolor*) females feed nestlings and remove fecal sacs more frequently than males [34]. However, in most bird species, males normally defend the nest more fiercely than females [34–37]. These sexual differences in parental investment may affect the relative reproductive value of the offspring for each sex, influencing the predation risk that each parent is willing to assume during the breeding period. Most previous studies evaluating the potential sexual differences in risk-taking behavior during the breeding period have exposed birds to the presence of a predator or a predator model in areas surrounding the nest [13,14,38]. Here, we examined differences in antipredatory behavior between female and male blue tit parents rearing nestlings in response to an increase in the risk of predation inside their nest-boxes, that may imply a greater risk of predation for parents than cues of a predator in the nest surroundings. To assess the risk taking assumed, we examined the latency to enter the nest-box for the first time, the provisioning rate and the time spent associated with sanitation activities. We expected that blue tit parents would detect predator chemical cues and exhibit antipredator behaviors, such as a reduction in nest sanitation activities or an increase in latency to enter the nest-box [11,14]. We did not expect that blue tit parents would modify their provisioning behavior, in accordance with previous results [11,14]. We also measured whether the number of begging nestlings differed in relation to the risk of predation to test whether nestlings responded to the chemical stimuli and whether their response might affect parental behavior. We hypothesized that sexual differences in reproductive investment as well as in confidence in parenthood may be related to the risk taking behavior of both parents. In blue tits, the relative value of the reproductive event is higher for females than for males [39, 40, 41] because parental investment is higher for females than for males during nest building, egg production and incubation, i.e. females invest more than males during the earlier stages of the breeding period. Moreover, confidence in parenthood is supposed to be higher for females because extrapair paternity is common (10–16% of nestlings are not fathered by the social father; [42–44], and, at least in Central Spain, nearly half of the nests contain at least one extra-pair nestling [43] while conspecific brood parasitism is low. Consequently, we expected that females would assume a greater risk than males if they find predator chemical cues inside the nest-box to maximize their reproductive success.

## Materials and methods

### Study species and area

The study was carried out during May and June 2017 in Pyrenean oak forests of Zaragoza (Alto Huerva-Sierra de Herrera) and Teruel (Sierra de Fonfría) provinces, in Spain (40°99′ N, 1°08′ W), where 250 nest-boxes were hung from tree branches 4 m above ground and had been available to the tits since 2011. Blue tits in our population lay a clutch of 9 eggs on average (range 7–12), incubated by the female for 12–16 days. Nestlings are fed by both parents and fledge at 16–23 days of age [32]. Nest-boxes (internal height × width × depth: 185 × 115 × 130 mm, bottom-to-hole height: 100 mm, hole diameter: 32 mm) were routinely checked to determine laying dates of the first egg, hatching dates and brood size.

### Experimental design and procedure

We assigned one treatment per nest when nestlings were 8 days old. The treatments were alternatively assigned to blue tit nests with similar number of nestlings and hatching dates. Some nests were later removed from the analyses because several cameras failed to record the behavior of birds, which resulted in unequal sample sizes between treatments. Forty-five blue tit nests were included in the experiment. Experimental treatments assigned to nests were: predatory mammal scent (mustelid; n = 19 nests), non-predatory mammal scent (odor control, rabbit (*Oryctolagus cuniculus*); n = 16) and water (odorless control; n = 10). There were no significant differences among treatments in hatching date (F_2,42_ = 0.07, p = 0.94), time of day when treatment was applied (F_2,42_ = 1.48, p = 0.24) and brood size (mean ± SE: predator: 7.58 ± 0.51; non-predator: 6.88 ± 0.43; control: 8.30 ± 0.50; F_2,42_ = 1.74, p = 0.19). However, differences among treatments in mean nestling mass approached significance but not in any systematic direction among treatments (mean ± SE: predator: 7.41 ± 0.24; non-predator: 7.97 ± 0.23; control: 6.98 ± 0.34; F_2,42_ = 3.15, p = 0.05).

We used ferrets (*Mustela putorius furo*) as the source of predatory mammal scent because its scent is similar to that of other mustelids [45] such as the European polecat (*Mustela putorius*), the weasel (*Mustela nivalis*) or the European pine marten (*Martes martes*), which are present in the study area and can prey upon adult and nestling birds [46,47]. We used rabbits (*Oryctolagus cuniculus*) as donors of non-predatory mammal scent because they are present in the study area but do not imply any risk of predation to birds. We used water as an odorless control stimulus to mimic the humidity of the experimental paper used in the other treatments.

We obtained predatory mammal and non-predatory mammal scents by placing clean absorbent papers inside the individual cages of two adult male ferrets and two adult male rabbits. We placed papers in each cage for 3 days before the experiment to ensure odor collection. The day of the experiment, we collected wet papers impregnated with recent cues of fresh urine. Papers soiled with feces were discarded. This method of odor collection has been used in previous studies [11,14,48–51]. Ferret scent is recognized by blue tits and great tits (*Parus major*) as a predatory threat [11,14,48,49] and rabbit scent has been used as an odorous control in other studies [14,50,52]. The two male scents within each treatment were alternately assigned to blue tit nests. The odorless control treatment was prepared by adding several drops of faucet water to clean pieces of absorbent paper. Water has been used as an odorless control stimulus in previous studies on the chemical ecology of birds [11,14,48,52].

Parent blue tits were captured with nest-box traps when tending 6-day-old nestlings. The sex of the adults was determined by the presence of a brood patch, present only in females. The first parent captured from each nest was marked with a small vertical white line made with Tipp-Ex© on the crown of the head to distinguish both parents in the video recordings. There were no significant differences between treatments in the ratio of males and females that were marked (*χ*^*2*^ = 0.17, d.f. = 2, p = 0.92). Routine body measurements and ringing were conducted before releasing birds. After capturing the parents, a dummy camera (and cable) was installed inside the nest-box, in the roof, to habituate birds to the presence of the real camera two days later.

When nestlings were 8 days old, we replaced the dummy camera by a real minicamera (Velleman® 6 IR LEDs mini color CMOS camera) connected by a cable to a recorder (DGD® mini C-DVR SD card recorder) and a 12-V battery located in a plastic container hidden under fallen leaves or vegetation below the nest-box. Two absorbent papers (11 x 13 cm) with the corresponding scent were hidden between the nest and the walls of the nest-box. Therefore, papers could not be visually detected or contacted by birds. However, birds could perceive the scent of experimental papers [11,48]. To examine the behavioral response of birds to the chemical cues of predators, the nest-box interior was filmed for 60 min immediately after placing the experimental stimulus. Video-recordings were conducted between 8:00 h and 16:00 h. The assignment of the different treatments was randomized relative to hour. After filming, we removed the absorbent papers from the nest-box and recorded body mass of all nestlings with an electronic balance (± 0.1 g) to control for potential differences between nests in nestling body mass that may affect the behavioral response of parents and nestlings to the treatments.

An observer, blind to treatments, analyzed video recordings and recorded for each parent: (a) latency to enter the nest-box for the first time (recorded as the time elapsed from the onset of filming until the parent entered the nest-box for the first time), (b) provisioning rate (number of provisioning events/hour; we considered a provisioning event when an adult visited the nest and fed the nestlings), (c) number of begging nestlings (mean number of nestlings that performed a begging response per adult visit; we considered a begging response when nestlings opened their beak for food solicitation), and (d) time spent in sanitation activities by females (i.e., other nest sanitation activities apart from fecal sac removal).

The experiment was conducted under a license issued by the Instituto Aragonés de Gestión Ambiental (INAGA/500201/24/2015/11696, INAGA/500201/24/2016/10017). No nest was abandoned during the course of the experiment. Exposure to the experimental odor treatments lasted only one hour and results of a previous study showed that exposure to predator chemical cues does not affect body condition of blue tit nestlings, even when cues are located inside nest-boxes for 5 days [11].

### Statistical analyses

Continuous variables that did not meet parametric assumptions were Box-Cox transformed to ensure normality. Transformed variables, as well as model residuals, followed a normal distribution. We used ANOVA to test whether brood size, hatching date and mean nestling body mass differed between treatments. A generalized linear model with binomial errors and a logit link function was used to analyse whether the ratio of males and females that were marked differed in relation to the treatment. We built general linear models with repeated measures to analyze whether there were differences between males and females (within-subject factor) and treatments (between-subject factor) in: (a) latency to enter, (b) provisioning rate, and (c) number of begging nestlings. We included the interaction between treatment and sex to examine whether the response to the different treatments differed between females and males. We controlled for variation in laying date, time of day, brood size and mean nestling mass between nests by including these variables as covariates in the analyses. In the provisioning rate analyses, we also controlled for variation in latency to enter by including these variables as covariates, because there were negative correlations between latency to enter and provisioning rate for females (Spearman’s correlation: r = − 0.74, p < 0.05), and for males (Spearman’s correlation: r = −0.82, p < 0.05). We also analyzed with a general linear model whether there were differences between treatments in (d) the time devoted to nest sanitation activities by females. We did not include males in this model because males never cleaned the nest. We controlled for variation in laying date, time of day, brood size and mean nestling mass between nests by including these variables as covariates in the analyses. We also controlled for variation in provisioning rate of females by including this variable as a covariate, because there was a positive correlation between time spent in sanitation activities and provisioning rate (Spearman’s correlation: r = 0.62, p < 0.05).

Non-significant terms were removed from the saturated models by a backward stepwise selection procedure in order to maximize multiple R^2^ of the models. Pairwise comparisons were planned using the Tukey HSD test. We corrected for multiple testing using the algorithm developed by Benjamini and Hochberg [53] to control the false discovery rate (FDR). This method is more suitable to ecological research than the less powerful and very conservative Bonferroni procedures (e.g. [54]). A prerequisite to wisely apply FDR or other multiple testing procedures is to define appropriate groups or families of hypotheses [53,54]. In our study, three families of hypotheses can be conservatively distinguished in relation to the behavioral response of parents and nestlings (latency to enter the nest-box, provisioning rate and number of begging nestlings): those concerning with the effect of sex (N = 3 tests, all P values ≥ 0.003 not significant after FDR control), those concerning with the effect of treatment (N = 3 tests, all P values not significant after FDR control), and those concerning interaction between treatment and sex (N = 3 tests, all P values not significant after FDR control). Significant results are indicated after correcting for multiple testing to control the false discovery rate. Statistical analyses were performed with STATISTICA 8.0.

## Results

### (a) Latency to enter

Latency to enter the nest-box after treatment presentation was shorter for females than for males (means in Table 1, F_1,42_ = 208.51, p = 0.003, p corrected for multiple testing to control the false discovery rate). However, there were no significant differences among treatments. The interaction between treatment and sex was not significant either. The latency to enter was not influenced by laying date, time of day, brood size or nestling body mass (saturated model in Table 2).

**Table 1.** Mean ± SE latency to enter the nest-box for the first time, provisioning rate, number of begging nestlings for male and female blue tits, and time devoted to nest sanitation activities by females in relation to experimental treatments: water (odorless control), non-predatory mammal scent (odorous control) and predatory mammal scent.

|  | Odorless control |  | Non-predatory mammal scent |  | Predatory mammal scent |  |
| --- | --- | --- | --- | --- | --- | --- |
|  | Female | Male | Female | Male | Female | Male |
| Latency to enter (s) | 1473.20 ± 384.91 | 1641.00 ± 433.54 | 1660.44 ± 283.01 | 2305.50 ± 287.00 | 1735.90 ± 285.61 | 2444.53 ± 265.06 |
| Provisioning rate | 9.10 ± 1.95 | 4.40 ± 1.31 | 6.38 ± 1.50 | 4.63 ± 1.40 | 6.16 ± 1.87 | 2.32 ± 0.66 |
| Number of begging nestlings | 4.40 ± 0.52 | 7.28 ± 4.48 | 2.39 ± 0.43 | 3.83 ± 1.88 | 3.32 ± 0.51 | 2.53 ± 0.60 |
| Time spent cleaning the nest (s) | 145.90 ± 41.80 | | 248.88 ± 118.33 | | 61.68 ± 30.62 | |

**Table 2.**
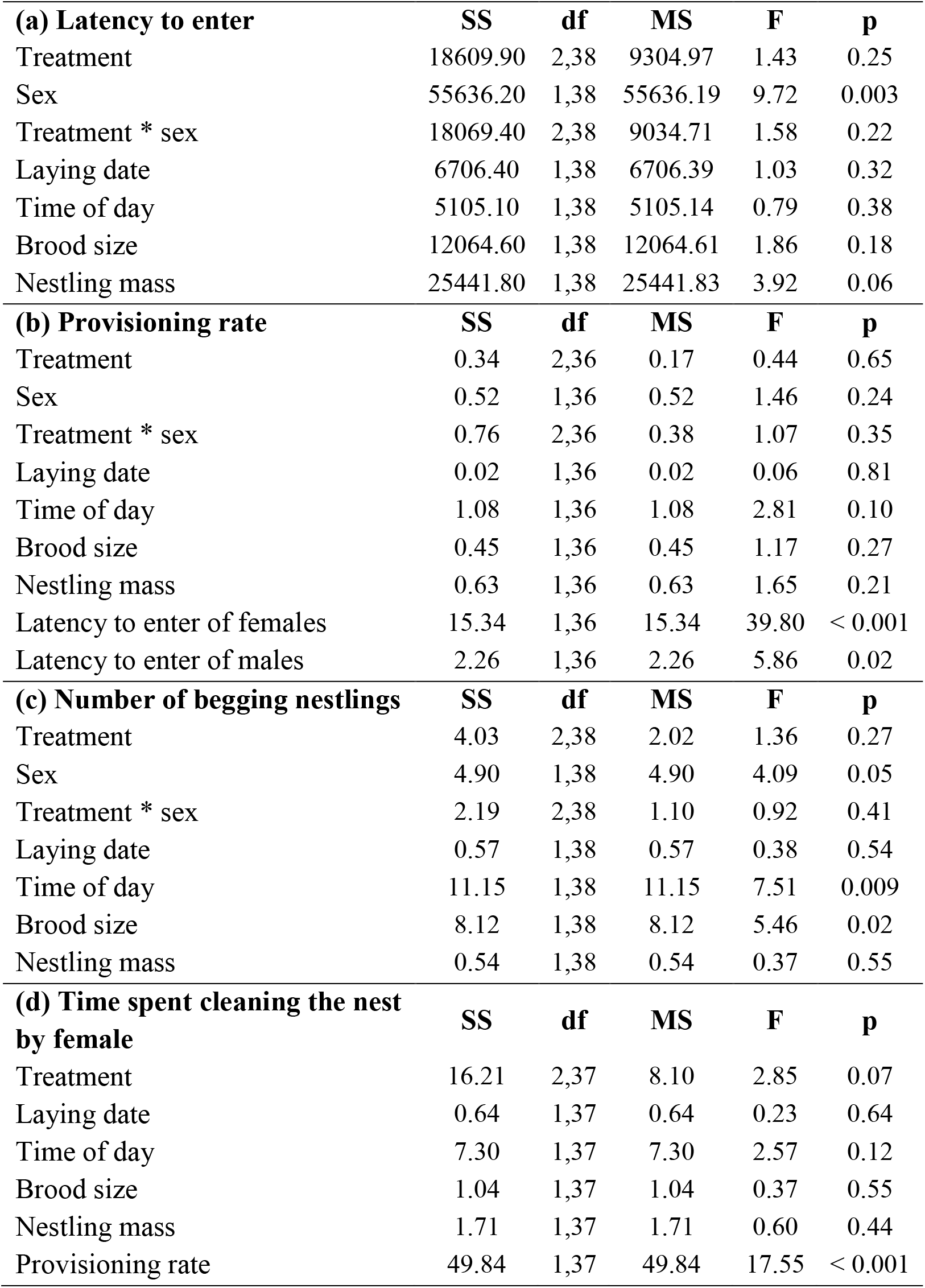
Saturated General Linear Models exploring the effects of treatment (control, non-predatory mammal, predatory mammal) and sex (females and males), and their interaction, on behavioral variables of blue tit parents feeding 8-day-old nestlings. Laying date, time of day, brood size and mean nestling mass were included as fixed covariables in all analyses.

|  |  |  |  |  |  |
| --- | --- | --- | --- | --- | --- |
| **(a) Latency to enter** | **SS** | **df** | **MS** | **F** | **p** |
| Treatment | 18609.90 | 2,38 | 9304.97 | 1.43 | 0.25 |
| Sex | 55636.20 | 1,38 | 55636.19 | 9.72 | 0.003 |
| Treatment * sex | 18069.40 | 2,38 | 9034.71 | 1.58 | 0.22 |
| Laying date | 6706.40 | 1,38 | 6706.39 | 1.03 | 0.32 |
| Time of day | 5105.10 | 1,38 | 5105.14 | 0.79 | 0.38 |
| Brood size | 12064.60 | 1,38 | 12064.61 | 1.86 | 0.18 |
| Nestling mass | 25441.80 | 1,38 | 25441.83 | 3.92 | 0.06 |
| **(b) Provisioning rate** | **SS** | **df** | **MS** | **F** | **p** |
| Treatment | 0.34 | 2,36 | 0.17 | 0.44 | 0.65 |
| Sex | 0.52 | 1,36 | 0.52 | 1.46 | 0.24 |
| Treatment * sex | 0.76 | 2,36 | 0.38 | 1.07 | 0.35 |
| Laying date | 0.02 | 1,36 | 0.02 | 0.06 | 0.81 |
| Time of day | 1.08 | 1,36 | 1.08 | 2.81 | 0.10 |
| Brood size | 0.45 | 1,36 | 0.45 | 1.17 | 0.27 |
| Nestling mass | 0.63 | 1,36 | 0.63 | 1.65 | 0.21 |
| Latency to enter of females | 15.34 | 1,36 | 15.34 | 39.80 | < 0.001 |
| Latency to enter of males | 2.26 | 1,36 | 2.26 | 5.86 | 0.02 |
| **(c) Number of begging nestlings** | **SS** | **df** | **MS** | **F** | **p** |
| Treatment | 4.03 | 2,38 | 2.02 | 1.36 | 0.27 |
| Sex | 4.90 | 1,38 | 4.90 | 4.09 | 0.05 |
| Treatment * sex | 2.19 | 2,38 | 1.10 | 0.92 | 0.41 |
| Laying date | 0.57 | 1,38 | 0.57 | 0.38 | 0.54 |
| Time of day | 11.15 | 1,38 | 11.15 | 7.51 | 0.009 |
| Brood size | 8.12 | 1,38 | 8.12 | 5.46 | 0.02 |
| Nestling mass | 0.54 | 1,38 | 0.54 | 0.37 | 0.55 |
| **(d) Time spent cleaning the nest by female** | **SS** | **df** | **MS** | **F** | **p** |
| Treatment | 16.21 | 2,37 | 8.10 | 2.85 | 0.07 |
| Laying date | 0.64 | 1,37 | 0.64 | 0.23 | 0.64 |
| Time of day | 7.30 | 1,37 | 7.30 | 2.57 | 0.12 |
| Brood size | 1.04 | 1,37 | 1.04 | 0.37 | 0.55 |
| Nestling mass | 1.71 | 1,37 | 1.71 | 0.60 | 0.44 |
| Provisioning rate | 49.84 | 1,37 | 49.84 | 17.55 | < 0.001 |

### (b) Provisioning rate

Provisioning rate of females was higher than that of males (means in Table 1, F_1,40_ = 28.69, p < 0.001). However, there were no significant differences among treatments. The interaction between treatment and sex was not significant either. Provisioning rate was not influenced by laying date, time of day, brood size or mean nestling mass. However, provisioning rate was negatively correlated with latency to enter in females (F_1,40_ = 51.74, p < 0.001) and in males (F_1,40_ = 10.55, p = 0.002) (saturated model in Table 2).

### (c) Number of begging nestlings

There were no significant differences in the number of begging nestlings between treatments and sexes, and the interaction between treatment and sex was not significant. The number of nestlings that performed a begging response did not differ in relation to laying date and mean nestling mass. However, the number of nestlings that performed a begging response per adult visit was negatively correlated with time of day (F_1,40_ = 7.20, p = 0.01) and positively correlated with brood size (F_1,40_ = 7.13, p = 0.01) (saturated model in Table 2).

### (d) Time devoted to nest sanitation activities by females

The time devoted to nest sanitation activities by females differed in relation to treatment (F_2,41_ = 3.81, p = 0.03, means in table 1, Fig 1). Females decreased the time devoted to nest sanitation activities when there were chemical cues of a predator inside the nest, although differences were only significant when comparing the control and the predator treatments (p = 0.04). The time devoted to nest sanitation activities by females was not influenced by laying date, time of day, brood size and mean nestling mass, but was positively correlated with provisioning rate (F_1,41_ = 28.01, p < 0.001) (saturated model in Table 2).

**Fig 1.**
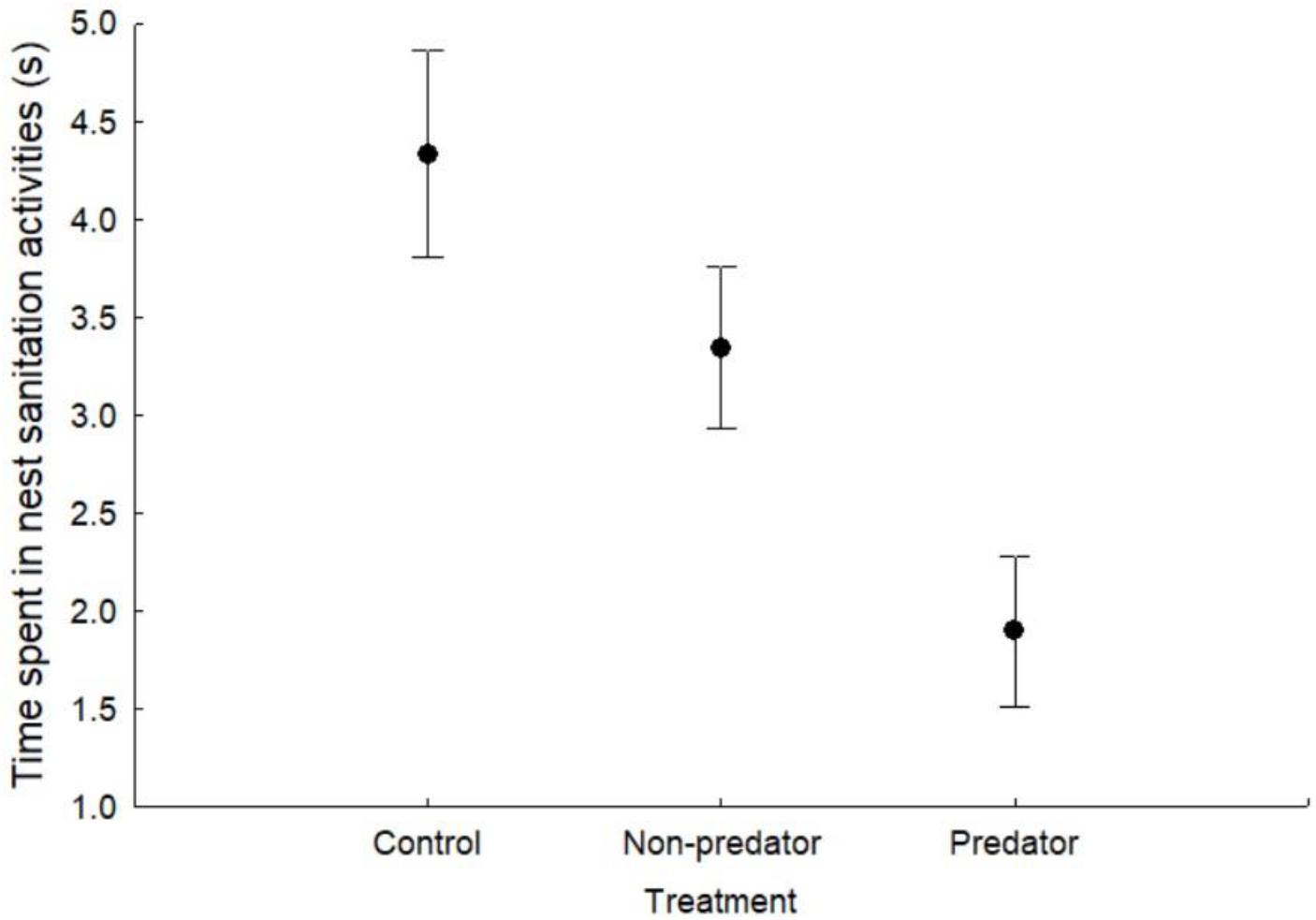
Mean ± SE time devoted to nest sanitation activities by blue tit females in relation to treatment: control, non-predatory mammal (rabbit), and predatory mammal (mustelid) chemical cues placed inside the nest-box. Figure shows Box-Cox transformed data.

## Discussion

Our results suggest that blue tits detected different chemical cues inside nest-boxes and used it to assess the level of predation risk. However, contrary to our expectations, we found that males and females did not differ in the risk assumed in response to perceived predation risk. Despite the reproductive value of the offspring should be higher for females than for males, we did not find that females were more prone to enter the nest-box than males. This result is in accordance with previous results that showed that blue tits and pied flycatchers (*Ficedula hypoleuca*) did not show sexual differences in the latency to enter in relation to perceived predation risk when exposed to stuffed models outside the nest cavity [13,52]. We also did not find differences in latency to enter the nest-box among treatments, in contrast to previous results in this species, where blue tits delayed their entry to the nest-box when there were predator chemical cues inside it. Differences in the results can be explained because in this study we captured adults to mark them whereas in the first study, adults were not captured before the experiment. The capture of adults two days before the experiment may have caused adults to be more cautious when entering the nest-box when they perceived any change due to experimental setting, and this may have masked any possible effect of treatment (see [55]). The higher latency to enter the nest-box in this study (1650 ± 175 s) compared to the previous study (342 ± 83 s, [11]) support this explanation.

Our results also show that there were no significant differences between females and males in provisioning rate in relation to predation risk, in accordance with previous studies in blue tits. For example, Mutzel and collaborators did not find sexual differences in nest defense behavior and provisioning rates during the breeding period [13]. Similarly, pied flycatchers reduced the parental provisioning rate when exposed to stuffed sparrowhawk (*Accipiter nisus*) models, but there were no sexual differences in this parental antipredator response [56]. Also, male and female blackbirds (*Turdus merula*) responded similarly to the risk of nest predation, simulated in this case with calls of an avian nest predator in the surroundings of the nest, by reducing the visits to the nest [18]. Although several studies have found that birds reduced provisioning rates when they perceived an increase in predation risk, in order to decrease the probability of predation [8–10,12,20,23], our results show that blue tits did not reduce provisioning rates, according to previous results in this species [11,14]. By maintaining provisioning rates, blue tit parents ensure nestling growth and increase fledging success [11]. Parental provisioning rate may be related to begging intensity, as begging behavior can signal the hunger level and the body condition of nestlings [57], so parents often adjust provisioning rate to begging intensity [58]. However, because begging behavior can also attract predators and increase predation risk [20,59], nestlings may reduce their begging behavior when they detect a predator inside the cavity. For example, Briskie and collaborators found that nestlings reduced the intensity or the number of begging calls to minimize the possibilities that a predator could locate the nest and reduce the risk of attracting predators [60]. However, our results show that blue tit nestlings did not modify their begging behavior when they were experimentally exposed to the chemical cues of a predator inside the nest cavity. The lack of responsiveness to predator scent in nestlings may not allow us to disentangle whether blue tit nestlings were able to recognize the predator scent, but the innate detection of predator scent has been previously found in a closely related species, the great tit [50]. Further experiments are needed to examine whether the ability to discriminate between threatening and non-threatening chemical cues is innate in blue tits, and whether the response varies with experience [61].

The results also show that blue tit females reduced nest sanitation activities when they detected predator chemical cues inside the nest-box. This is in accordance with results of a previous study where visual or chemical cues of predators were placed in the vicinity of the nest, instead of inside the nest-box, where this increase in the risk of predation elicited a reduction in nest sanitation activities of parents [14]. By decreasing these activities that are not essential for nestling survival, females may minimize the time exposed to the predator inside the nest cavity as an antipredatory strategy to minimize their own risk of predation. However, this antipredatory behavior may entail indirect costs in terms of increased ectoparasite loads in the nest, with detrimental consequences for the fitness of both adult and nestling birds [19].

In conclusion, this study adds to previous evidence that birds detect the chemical cues of predators in the nesting cavity ([11,48,50]; but see [62,63]). Despite we simulated an elevated risk of predation by adding predator chemical cues inside the nest cavity, our results showed that the behavioral response of parents was no sex-dependent, in accordance with previous studies where predation risk was perceived in the nest surroundings. Sexual differences in the risk assumed may be related to differential parental investment and reproductive value of offspring for males and females. These factors may change over time during a reproductive event. In blue tits, females invest more time and energy during the first stages of reproduction (nest building, egg laying and incubation), but during nestling stage, both sexes devote similar time and energy in provisioning nestlings, and males invest more in nest defense [39]. Then, sexual differences in reproductive investment might be reduced in the course of a reproductive event, and this may explain the lack of sexual differences we observed in risk taking behavior. Further experimental studies are needed during the first stages of reproduction to disentangle how female and male parents respond to perceived predation risk in relation to their current reproductive investment.

## Acknowledgements

We thank two anonymous referees for their helpful comments. I. Saavedra was supported by a FPI grant from the Spanish Ministry of Economy and Competitiveness. L. Amo and G. Tomás were supported by the Ramón y Cajal programme. G. Tomás was supported by grants CGL2017-89063-P and PID2020-117429GB-C21.

